# Gene ORGANizer: Linking Genes to the Organs They Affect

**DOI:** 10.1101/106948

**Authors:** David Gokhman, Guy Kelman, Adir Amartely, Guy Gershon, Shira Tsur, Liran Carmel

**Author notes:** These authors contributed equally to the work (David Gokhman, Guy Kelman).

## Abstract

One of the biggest challenges in studying how genes work is understanding their effect on the physiology and anatomy of the body. Existing tools try to address this using indirect features, such as expression levels and biochemical pathways. Here, we present Gene ORGANizer (geneorganizer.huji.ac.il), a phenotype-based tool that directly links human genes to the body parts they affect. It is built upon an exhaustive curated database that links more than 7,000 genes to ∼150 anatomical parts using >150,000 gene-organ associations. The tool offers user-friendly platforms to analyze the anatomical effects of individual genes, and identify trends within groups of genes. We demonstrate how Gene ORGANizer can be used to make new discoveries, showing that chromosome X is enriched with genes affecting facial features, that positive selection targets genes with more constrained phenotypic effects, and more. We expect Gene ORGANizer to be useful in a variety of evolutionary, medical and molecular studies aimed at understanding the phenotypic effects of genes.

---

Many high-throughput methods such as whole-genome sequencing, expression microarrays, RNA-seq, and whole-genome methylation mapping produce genome-wide data whose analyses produce long lists of genes of interest. These lists typically include genes that share a certain trait such as being bound by the same transcription factor, being differentially methylated between two samples, having high conservation levels, or being differentially expressed following a treatment. Such lists have become a common product of biological research, but understanding how they affect the biology of an organism at the physiological and anatomical level remains a challenging task^1^.

Dozens of tools have been developed to address this challenge, providing researchers with powerful means to tease out biological processes and functions that are associated with the genes they investigate^1–3^. For example, a popular tool is DAVID^2,4^, where genes can be analyzed for shared Gene Ontology (GO) terms, disease associations, expression patterns and biochemical pathways. The strategy adopted by many of these tools, e.g., Human Phenotype Ontology (HPO)^5^, DisGeNet^6^, PhenGenl^7^, PhenomicDB^8^ and Organ System Heterogeneity DB^9^, is to focus on the phenotypic effects of genes. Thus, these tools usually harbor databases (DBs) for gene-phenotype associations. However, genes in these DBs are linked either to diseases (e.g., ‘primary ciliary dyskinesia’), or to the phenotypes of a disease (e.g., ‘peripheral traction retinal detachment’), but not directly to organs (e.g., ‘eye’). Tools such as OMIM^10^, Organ System Heterogeneity DB^9^ and BRITE^11^ do offer some direct links between genes and organs, but include only a limited number of organs and systems (33 in OMIM, 26 in Organ System Heterogeneity DB, 12 in BRITE), and lack platforms to efficiently mine and analyze these data.

Another approach for linking genes to body parts is based on expression rather than phenotype, where mRNA levels are used to determine in which tissues and cell types genes are active. For example, Expression Atlas^12^ is a tool allowing analysis of gene expression in different cell types, diseases and developmental stages, based on comprehensive RNA-seq and microarray data. While very useful in many cases, expression-based analysis is an indirect approach that suffers from a number of drawbacks. First, the repertoire of expression datasets is limited, with a strong bias towards certain organs and tissues (e.g., brain, blood and skin), whereas many other body parts are rare or completely absent (e.g., bone, face, larynx, urethra, teeth, fingers, and spinal cord). Second, samples used for expression analyses are usually obtained from specific developmental stages, taken post-mortem, and extracted from particular parts of the organ. Thus, the data collected rarely capture the entire temporal and structural variation of organs. Third, expression analyses generally focus on specific cell types or tissues (e.g., cardiomyocytes), rather than on whole organs (e.g., heart), systems (e.g., the cardiovascular), or anatomical regions (e.g., the thorax), hence providing partial or skewed information on how whole organs are affected. Finally, gene expression does not directly translate into an observable phenotype. This limited correspondence between expression and phenotype stems from several reasons: (a) The correlation between mRNA levels and protein levels is generally low, reported to be less than 0.5^13–16^. (b) Expression assays, especially if done in low coverage, might miss lowly expressed genes. However, these genes tend to be more medically relevant and underlie organ-specific phenotypes^17^. (c) The activity of a gene is not necessarily limited to the tissue in which it is expressed. For example, expression of a gene in the endocrine system would often have phenotypic consequences in other tissues, due to its secretory function.

Thus, despite the plethora of tools designed for the analysis of gene lists, direct association of genes to body parts is largely unavailable. Today, researchers who seek to link genes to the organs they affect are left with two main options: either to use gene expression DBs, which do not provide a direct phenotype-based association, or to conduct a manual review of the literature and free text DBs such as OMIM^10^, Gene Cards^18^ and GenBank^19^, which are not constructed for gene list analyses.

Gene ORGANizer was developed to fill this gap. We have constructed a comprehensive fully curated DB, consisting of more than 150,000 gene-body part associations, and covering over 7,000 human genes. The body parts are divided into four levels of hierarchy: body systems (e.g., cardiovascular, hereinafter *systems*), anatomical regions (e.g., thorax, hereinafter *regions*), organs (e.g., heart) and germ layers (e.g., mesoderm). On top of this DB, we have created a web platform that allows users to browse for a specific gene, as well as to analyze gene lists in order to test whether they are enriched or depleted with certain body parts.

## Results

### Backend database

In non-human organisms phenotypes can be directly observed using various genetic manipulations such as knockout or knockdown. In humans, however, the principal way to associate genes to phenotypes is through observed diseases. To construct the Gene ORGANizer DB, we used two of the largest DBs for gene-disease and gene-phenotype associations in human: Human Phenotype Ontology (HPO)^5^ and DisGeNET^6^. HPO integrates data from three highly-curated sources: OMIM^10^, Orphanet^20^ and DECIPHER^21^. DisGeNET integrates data from UniProt^22^, The Comparative Toxicogenomics Database (CTD)^23^, and ClinVar^18^, as well as from non-human sources, such as CTD mouse^23^, CTD rat^23^, The Mouse Genome Database (MGD)^24^ and The Rat Genome Database (RGD)^25^. DisGeNET also includes annotations based on literature text mining, which we do not use for Gene ORGANizer, as they are not curated. Together, these DBs link 7,132 human genes to diseases and phenotypes (see online Methods).

We have built our tool based on the entire HPO DB and the curated portion of DisGeNET, which together comprise over 150,000 gene-phenotype and gene-disease associations. We developed a protocol to translate these data into associations between genes and the anatomical parts in which the phenotype is observed (Fig. 1). For example, one of the phenotypes caused by mutations in the *HOXA2* gene is microtia – the underdevelopment of the outer ear (OMIM ID: 612290)^26^. We have used this association to link *HOXA2* to the following body parts: the outer ear, the ear, the head, the integumentary system, the head and neck region and the ectoderm germ layer (see online Methods for a complete description of the annotation protocol). Overall, we have linked genes to 146 body parts, divided into four anatomical hierarchies: (a) three germ layers (endoderm, mesoderm and ectoderm); (b) six regions (head and neck, thorax, abdomen, pelvis, limbs and non-specific); (c) twelve systems (digestive, nervous, reproductive, endocrine, skeletal muscle, skeleton, lymphatic, cardiovascular, immune, urinary, respiratory and integumentary); and (d) 125 organs and sub-organs (Supplementary Table S1).

**Fig. 1.**
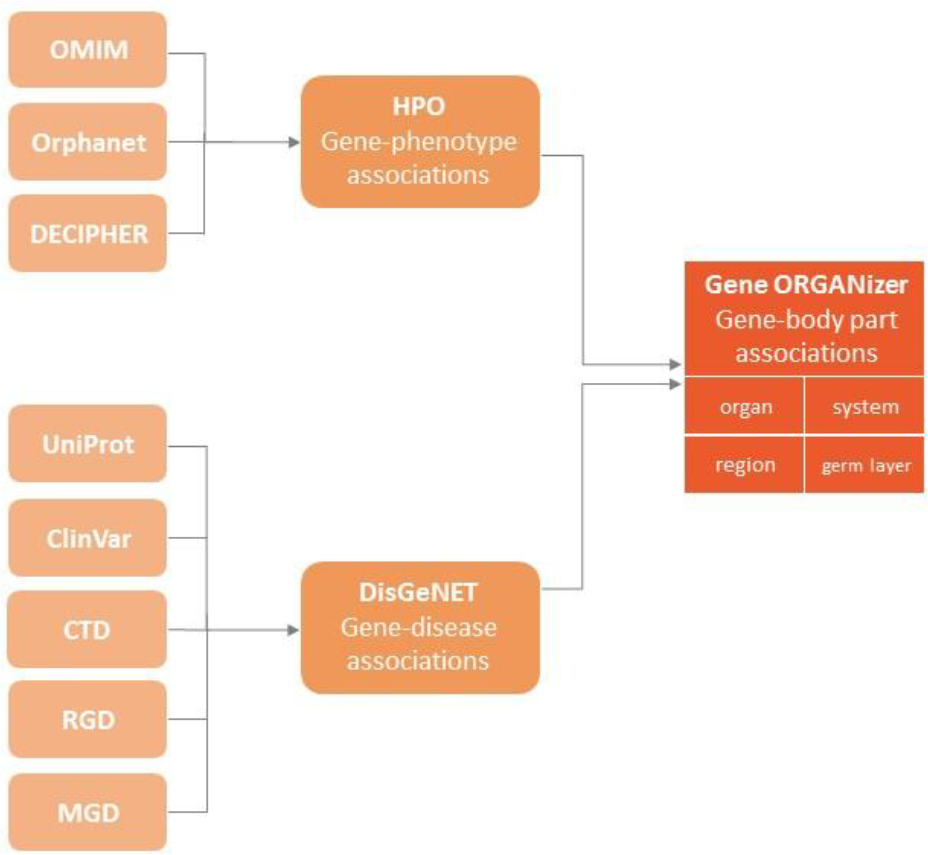
Sources of the Gene ORGANizer database. Sources of associations that comprise the Gene ORGANizer DB. Associations in Gene ORGANizer are divided into four levels of hierarchy: organ (e.g., stomach), system (e.g., digestive), region (e.g., abdomen) and germ layer (e.g., endoderm).

### Using Gene ORGANizer

Gene ORGANizer was designed to provide researchers with the ability to analyze the phenotypic effects of genes and to understand the shared impact of groups of genes. The tool consists of two platforms: *Browse* and *ORGANize*. *Browse* allows users to see all of the body parts affected by a single gene of interest. *ORGANize* is designed to test which body parts, if at all, are over- or under-represented in a gene list. In both platforms, the user can base the analysis on either the *typical* phenotypes associated with a gene (defined as those that appear in more than 50% of sick individuals), or on its *typical+non-typical* phenotypes (i.e., any frequency). Additionally, the user can choose between *confident* associations (i.e., inferred from data on humans), and *confident+tentative* ones (inferred also from additional data on mouse and rat).

The output in both *Browse* and *ORGANize* comes in two forms: a color-coded body map and a table. The table contains all information whereas the body map visualizes most of it (125 out of the 146 body parts). Non-localized body parts (e.g., blood) or very small parts (e.g., sweat gland) do not appear in the body map, and are represented only in the table. In the *Browse* option, the table and body map simply present the body parts that are phenotypically affected by the gene of interest, colored by the type of association (*confident* or *tentative; typical* or *non-typical*). Hovering over a body part in the table allows the user to see the phenotypes and diseases that are behind the gene-body part association. In the *ORGANize* option, the body map represents an interactive heat map, where significantly enriched or depleted body parts are colored based on the level of their enrichment or depletion. Non-significant body parts remain in their original grey color.

The enrichment and depletion tests within a gene list are carried out against a list of background genes. By default, the background consists of all genes that are linked to body parts in our DB. This background assures that even if certain anatomical parts are over-represented in the ontology (because some phenotypes are easier to detect, or some diseases are more studied), it would not bias the results^2^. Gene ORGANizer also allows users to enter their own background list. User-specified backgrounds are useful in cases where the initial pool of genes from which the gene list was derived contains an inherent bias. For example, in a list of genes that were found to be differentially regulated based on a microarray experiment, the background should comprise only genes that are represented on that microarray.

### Controlling for potential biases

To investigate potential biases in our DB, we ran Gene ORGANizer on random lists of 100, 500 and 1,000 genes, and tested how many significantly enriched or depleted body parts are reported for different types of associations – *confident*, *confident+tentative*, *typical* and *typical+non-typical*. We repeated this procedure 1,000 times and found that significantly enriched/depleted body parts were rarely observed. For example, for lists of 100 genes, only 0.5% of the *confident typical+non typical* iterations returned significant organs (FDR < 0.05), 4.2% for 500 genes and 3.8% for 1000 genes (Supplementary Table S2).

To further assess the level of accuracy in of our DB, we compared Gene ORGANizer to the OMIM organ annotations, which links disease to 33 of our 125 organs^10^. Comparing the two, we found that less than 1% of our annotations were not in accordance with OMIM’s.^11^

As a positive control we used housekeeping genes, which are genes that participate in basic cellular functions and are thus ubiquitously active and affect many anatomical parts^27^. On average, each housekeeping gene is expected to be linked to more organs than in the genomic background. In this case, Gene ORGANizer will produce substantially more enriched body parts than expected by chance. We ran Gene ORGANizer on 3,804 housekeeping genes^27^ and reassuringly, found that most systems (7 out of 12) and regions (5 out of 6) were significantly enriched, as well as 32 organs (Supplementary Table S3). Such high numbers of significant body parts are rarely observed at random (*P* = 0.001 for systems, *P* = 0.015 for regions, *P* = 0.003 for organs, randomization test of 3,804 genes).

As another positive control, we extracted from the Kyoto Encyclopedia of Genes and Genomes (KEGG)^11^ genes that are part of biochemical pathways linked to specific body systems. We did this for all body systems represented in KEGG, namely the circulatory, immune, endocrine, digestive, and nervous systems, and demonstrated how in each case, Gene ORGANizer identified the relevant body parts as significantly enriched (Supplementary Table S4). Within the genes in KEGG that are associated with the circulatory system, Gene ORGANizer identified that the most enriched organs are the heart valve (x2.17, FDR = 2·10^−7^), red blood cells (x1.69, FDR = 0.009) and the heart (x1.50, FDR = 5·10^−4^, Supplementary Fig. S1). Within immune-related genes, the most enriched systems were the lymphatic (x2.78, FDR < 10^−15^) and immune (x1.75, FDR = 8·10^−11^) and the most enriched organs were the sinuses (x5.14, FDR = 5·10^−8^), lymph nodes (x4.89, FDR < 10^−15^), and bone marrow (x4.08, FDR < 10^−15^, Fig. 2). The sinuses probably appear in this list due to the elevated activity of lymphocytes within them, and the systemic link between the mucosal immune system and susceptibility to infections^28^. Interestingly, additional characteristics of the immune system can be detected in these results. For example, the brain is significantly depleted, corresponding to the lack of lymphatic drainage system in the brain. However, the meninges is found to be significantly enriched, in accordance with the recent discovery that some lymphatic vasculature exists in the central nervous system in the form of lymphatic vessels in the tissues that surround the brain^29^. Within endocrine-related genes, the endocrine system was the most enriched (x1.58, FDR = 3·10^−4^). See Online Methods for additional validations.

**Fig. 2.**
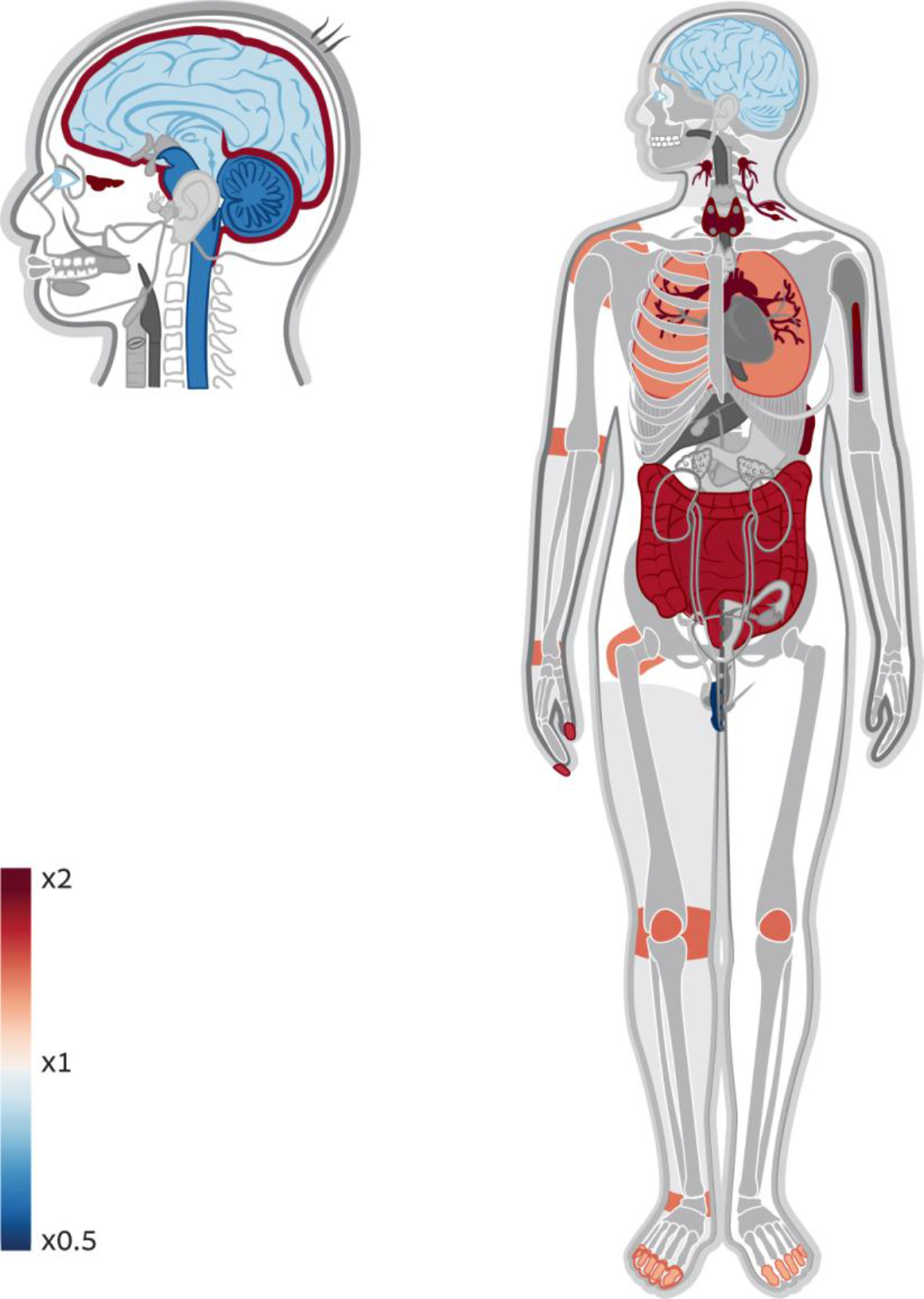
Gene ORGANizer detects enrichment of immune-related organs within immune-related genes. A body and head map of enrichment and depletion of organs across immune-related genes. As a positive control, we extracted from the Kyoto Encyclopedia of Genes and Genomes (KEGG)^11^ genes that are associated with specific systems. Genes that are involved in immune response were run in *ORGANize* and the most enriched body parts were those that are associated with immune response.

### Chromosome X is enriched with genes affecting facial features

Sex chromosomes have always been of special interest because of their distinctive evolutionary history and means of inheritance, which result in unique selection regime and disease manifestation^30–35^. The high occurrence of mental disorders in males drove researchers to look into chromosome X and investigate its link to the brain. Indeed, manual inspection of the OMIM DB has shown that chromosome X has more than 3-fold enrichment in genes associated with mental retardation, raising the hypothesis that there is an over-representation of brain-related genes on chromosome X^34^. Other studies have shown that chromosome X is enriched with reproduction-related genes, and in particular with genes that are expressed in the testes^35^. As only one body system was investigated in each of these studies, it was impossible to put these findings in a larger context of the entire body and see how these enrichments scale up compared to other body parts, and if they are unique. Using Gene ORGANizer, not only do we validate the enrichment of brain- and reproduction-related genes within chromosome X, but interestingly, we observe a stronger trend that could not have been detected with current tools and DBs. The brain and testes are only two out of 45 organs that are significantly enriched within this chromosome. Almost half of them, including the most enriched ones, are parts of the face, (e.g., the mouth, cheeks, lips, chin, teeth, forehead, nose, hair, jaws and outer ear, FDR < 0.05, Fig. 3). In fact, aside from the eyes, all facial parts are significantly enriched within X-linked genes. We also show that it is not only testes-related genes that are enriched within chromosome X, but most organs of the urogenital system. Finally, we detect over-representation of many parts of the skeletal system, including the rib cage, pelvis, joints, limb extremities, spinal column and skull.

**Fig. 3.**
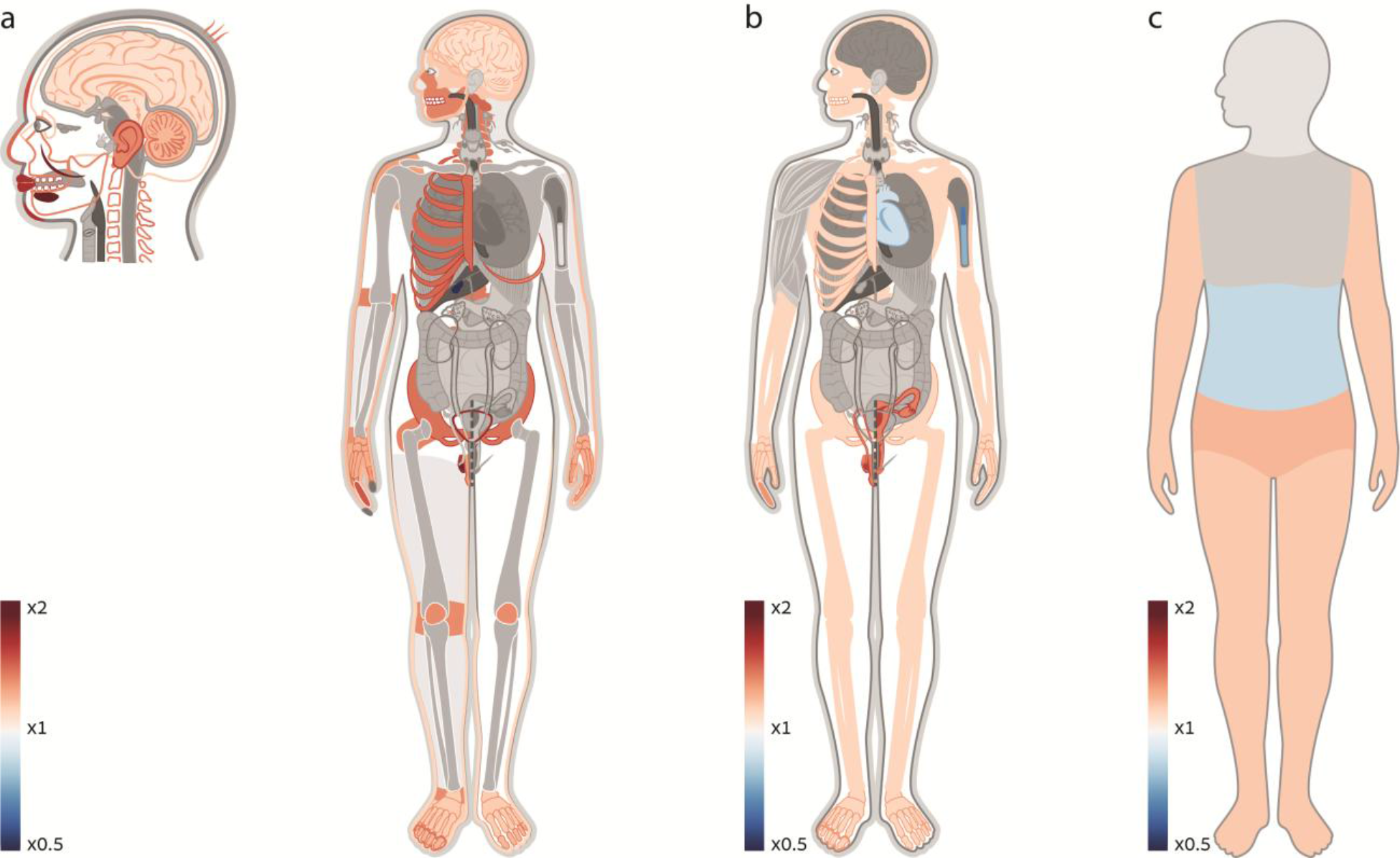
Genes affecting the face, the brain, and the urogenital and skeletal systems are over-represented on chromosome X. **a.** A heat map of enriched and depleted organs within X-linked genes. Gene ORGANizer detects significant enrichment of the brain and testes within these genes, confirming previous claims. A more pronounced trend is the over-representation of different facial features, including all parts of the face except the eyes. Many parts of the urogenital and skeletal systems are enriched as well. **b.** A heat map of enriched and depleted systems within X-linked genes. The reproductive and the skeletal systems are significantly enriched (x1.38 and x1.12, FDR = 3·10^−5^ and 0.022, respectively). The immune and the cardiovascular systems are significantly depleted (x0.74 and x0.87, FDR = 0.002 and 0.032, respectively). **c**. A heat map of enriched and depleted body regions within X-linked genes. The regions of the pelvis and limbs are significantly over-represented (x1.22 and x1.16, FDR = 5·10^−4^ and 0.003, respectively). The abdominal region is significantly depleted (x0.84, FDR = 0.008).

As a negative control, we applied Gene ORGANizer to chromosome 16, which resembles chromosome X in both size and number of genes. We found that the genes on chromosome 16 are not enriched with any body part (Supplementary Table S5). More generally, repeating the analysis for all other autosomal chromosomes revealed that their genes rarely show any significant association with specific body parts. The only chromosomes that showed any over-representation were chromosomes 9, 14 and 17, albeit to a much lesser extent compared to chromosome X, both in the number of enriched body parts and in the levels of enrichment (Supplementary Table S5). This suggests that chromosome X likely experiences a unique regime of selection leading to preferential representation of genes that affect the brain, the urogenital and skeletal systems, and above all - facial features.

A possible explanation for these observations is that being hemizygous in males, genes on chromosome X experience stronger and sex-specific selection compared to autosomal genes. This is because a newly emerged recessive allele on chromosome X will be expressed in males, but not in females. With this process in mind, Rice suggested in 1984 that genes on chromosome X play an important role in sexually dimorphic traits and in sexual selection^31^. In fact, based on Rice’s hypothesis, it is predicted that with time, sexually selected and sexually dimorphic genes will preferentially move, through chromosomal translocation, to chromosome X. Alternatively, this hypothesis predicts that X-linked genes will evolve sexually dimorphic function, and that they will be sexually selected for more often^31^. Indeed, it was shown later that chromosome X is highly enriched for genes that control sexually selected and sexually dimorphic traits^32,36^. Therefore, a possible explanation for our observations is that some of these organs are targets of sexual selection, and that their sexually dimorphic nature (such as in the case of the face, a classic sexually divergent^37,38^ and sexually selected organ^39^), was evolutionary advantageous.

These results emphasize the importance of Gene ORGANizer as a tool to investigate gene function outside the scope of gene expression data. Expression databases rarely provide information for body parts such as the face, and thus, they are restricted in the range of anatomical parts for which they can provide inference. This could explain how the most pronounced trend on chromosome X has not been detected to date.

### Imprinted genes tend to affect the same organs

Imprinted genes are genes that are transcribed only from one of the chromosomes – either the maternal or the paternal. This asymmetric silencing is achieved through DNA methylation of one of the alleles. This phenomenon evolved independently in plants and mammals, and its evolutionary role is still debated^40^. Aberrant imprinting, where both or none of the alleles are transcribed, results in a variety of abnormalities. Previous studies have shown that human imprinted genes within the same locus show similar temporal patterns of expression^40^. Concerted upregulation of imprinted genes from different loci has been identified as well^40^. Furthermore, imprinted genes have been shown to participate in similar biochemical pathways^40^. These observations suggest an intricate network of co-regulation of imprinted genes. However, the extent to which this phenomenon affects specific organs, and its phenotypic consequences are still to be determined^40^. To test this, we ran a list of 37 high-confidence imprinted genes^41^ in Gene ORGANizer. We used only *typical* annotations in order to examine only the most common effects of these genes. We found that the endocrine system is the most enriched system within imprinted genes, with an over-representation of x3.21 (FDR = 0.018, Supplementary Table S6). This suggests that much of the reported role of imprinted genes in the regulation of development and growth^40^ is executed through the endocrine system. Importantly, we show that organs previously hypothesized to be particularly influenced by imprinted genes (e.g., the brain^42^ and reproductive organs^43^) are not significantly enriched within these genes, compared to the rest of the genome. This emphasizes the importance of Gene ORGANizer as a tool that enables researchers to analyze associations with organs in a genome-wide context.

### Positively selected genes in hominids affect less organs

In order to understand natural selection in a wide context, it is crucial to examine its dynamics across many species. A recent study investigated patterns of natural selection across all extant *Hominidae* species (great apes, including humans)^44^. This study identified hundreds of genes that likely went through positive selection in each lineage. Although most signatures of positive selection are species-specific, we found shared phenotypic effects within these genes. Taking together the top 200 genes with the strongest signs of positive selection in each lineage (1581 unique genes in total), we found that 26 organs and 3 systems are significantly depleted (Supplementary Fig. S2, Supplementary Table S7). The only organs that show a limited degree of enrichment (albeit not significant) are related to the nervous system, in accordance with the GO annotation-based analyses in the original study^44^. Such across-the-board depletion suggests a more general possibility: these genes tend to affect less organs than expected by chance. Indeed, we found that positively selected genes along hominid lineages affect on average ∼5 organs less than random genes (29.4 compared to 34.5, *P* = 0.006, randomization test). This is also supported by the observation that some of the most depleted organs have ubiquitous functions that affect many aspects of the physiology (e.g., the parathyroid, hypothalamus, thymus, and thyroid). These results suggest an intriguing possibility that positive selection tends to occur in genes with narrower and more organ-specific functions.

## Discussion

Although the Gene ORGANizer DB is based mostly on human phenotypes, these associations probably hold to a large extent in other species. By converting a list of gene IDs from a non-human organism to human gene IDs, or by entering gene symbols, which are mostly shared between species, researchers can use our tool to analyze gene function in non-human organisms. In order to test this, we ran in Gene ORGANizer a list of 117 genes that show signals of convergent evolution in bats and dolphins^45^. As these mammals independently evolved echolocation, we expected this list to be enriched for genes that affect echolocation-related organs, such as the inner and middle ear. Indeed, we find these organs to be significantly enriched (x3.60 and x2.42, *P* = 0.001 and 0.003, respectively). We also ran Gene ORGANizer on genes where signals of positive selection were detected in the gibbon genome^46^. Possibly reflecting the exceptional arboreal locomotion of gibbons and their unique skeletal structure, we show how all subcranial bones and joints are significantly over-represented. We also find enrichment in organs related to the digestive, cardiovascular and nervous system (FDR < 0.05, Supplementary Table S8). When researchers first came to analyze the gibbon genome and assign such genomic regions with functional meaning, they were limited to the use of tools that were mainly designed for molecular- and pathway-level analyses^46^. Using Gene ORGANizer, we show how higher level anatomical analysis could be easily performed, and how this could provide researchers with novel results in both human and non-human genomes.

The annotation behind Gene ORGANizer produced a binary matrix of associations (see *downloads* tab on geneorganizer.huji.ac.il). This matrix reveals a system of links between genes and organs, and could be used to study the genetic interactions between organs. For example, the DB can be used to build a graph whose vertices are organs, where the strength of an edge between two organs is determined by the number of genes that regulate both organs. Such an analysis could shed light on genetic co-regulation of different organs, and help explain co-occurrence of various phenotypes^47,48^ at the macro and micro levels.

We presented here the Gene ORGANizer DB and tool for the phenotypic analyses of gene-organ associations. We trust that Gene ORGANizer could be useful in nearly any genome-wide study where questions related to anatomy are raised, whether from an evolutionary, medical or biochemical perspective.

## Acknowledgements

We thank The Human Phenotype Ontology Consortium (HPO) and DisGeNet for their comprehensive data, and Shiran Bar, Ido Sagi, Sagiv Shifman, Michal Linial and Benny Yakir for their advice and ideas. The work was supported by the Israel Science Foundation FIRST individual grant (ISF 1430/13).

## Online Methods

### Database annotation

We developed two pipelines to create gene-organ associations. The first was designed to use the information within the Human Phenotype Ontology (HPO)^5^ database (DB). HPO translates gene-disease associations from OMIM^10^, Orphanet^20^ and DECIPHER^21^ into gene-phenotype associations. For example, mutations in the *PROC* gene are known to be behind the *thrombophilia due to protein C deficiency (THPH3)* disease (OMIM ID: 176860). In HPO, *PROC* is linked to the known phenotypes of the disease, such as *warfarin-induced skin necrosis* (HP:0001038). HPO includes 153,576 such gene-phenotype associations for 3,526 genes (build 110, Jan 25, 2016). Based on these associations, and using four medical resources that harbor extensive information relating to phenotypes (MedScape^49^, OMIM^10^, Orphanet^20^ and UpToDate^50^), we extracted the body parts that are associated with each phenotype and translated gene-phenotype associations into gene-body part associations.

The second pipeline was designed to use information within the DisGeNET DB^6^, which harbors gene-disease associations that are ranked according to their level of curation: (1) *curated data* from CTD human^23^, Orphanet^20^, ClinVar^51^, GWAS catalog^52^ and UniProt^53^; (2) *predicted data* from rodents gene-disease associations, i.e., CTD mouse^23^, CTD rat^23^, MGD^24^ and RGD^25^ that were translated into human gene-disease associations; and (3) *literature-based* text-mining algorithms. For Gene ORGANizer we used only the curated and predicted data, and left out literature-based data. Importantly, DisGeNET links genes to diseases, not to phenotypes. To link the genes to body parts, we mapped diseases to phenotypes using the HPO DB, which includes gene-disease-phenotype associations, and proceeded as described above. Diseases in DisGeNET that do not appear in HPO were associated directly with the body parts that they affect (e.g., genes that were associated with *Thanatophoric dysplasia, type 1,* which is characterized, among other signs, by abnormality of the femur, were linked to the organs: *femur, thigh* and *upper limb,* to the *skeletal* system*, to the mesoderm* germ layer and to the *limbs region*). Here too, the associations were based on medical data from MedScape^49^, OMIM^10^, Orphanet^20^ and UpToDate^50^.

Each body part belongs to one of four hierarchies. In total, our DB includes 125 organs (Supplementary Table S1), twelve systems (*nervous, endocrine, lymphatic, cardiovascular, skeletal muscle, skeleton, integumentary, immune, reproductive, respiratory, digestive* and *urinary*), six regions (*head and neck, thorax, abdomen, pelvis, limbs* and *general*) and three germ layers (*endoderm, mesoderm* and *ectoderm*). Phenotypes were linked to multiple body parts in a nested structure. For example, the *ARID1A* gene, which is associated with the HPO phenotype *Absent fifth fingernail* (HP:0200104) was linked to *upper limb*, to the *limbs* region, to the *integumentary* system, and to the *ectoderm* germ layer. The full list of annotations can be downloaded from the *Downloads* tab on geneorganizer.huji.ac.il. See Supplementary Table S9 for the entire nesting structure).

The group of organs includes both classic organs (e.g., heart, kidney, etc.), and body parts that do not fall under the classic definition of an organ (i.e., a set of tissues, grouped together into a distinct structure and performing a specialized task), but appear in many phenotypes (e.g., *eyelid, cheek, head, outer ear, ankle*) The decision whether to include a body part in the list of organs was taken based on the number of phenotypes that affect this body part. If an organ was associated with less than 10 phenotypes, it was not granted a distinctive term, but was rather joined to the organ to which it belongs. For example, the jejunum was too rare to be counted as an independent organ, and therefore phenotypes that affect the jejunum were associated with the *small intestine*. On the other end, if a body part that is not a ‘classic organ’ was linked to many phenotypes, it was given a distinct term (e.g., *eyelid, sinus*).

HPO labels phenotypes that are observed in more than half of the disease cases as ‘typical’. Gene ORGANizer allows users to choose whether to analyze their list based on typical gene-body parts associations, or on all the associations. Additionally, HPO includes a hierarchal annotation system. For example, a gene that is linked to the phenotype *warfarin-induced skin necrosis* (HP:0001038), will also be linked to *Dermatological manifestations of systematic disorders,* to *Generalized abnormality of the skin*, to *Abnormality of the skin* and to *Abnormality of the integument*. For Gene ORGANizer, we used only the final and most specific level of annotation. Finally, ambiguous phenotypes and phenotypes that could not be linked to specific organs were discarded (e.g., *autosomal inheritance, pain, difficulty walking, exercise intolerance, asymmetric growth*). The pipeline that was based on DisGeNET could not be categorized into *typical* and *non-typical*, as this DB does not contain phenotype associations and their relative prevalence. Therefore, when converting gene-disease associations into gene-body part associations, only the main body parts affected by a disease were linked to the gene, and all gene-body part associations are tagged as *typical*.

Users can enter gene lists using any mixture of the following gene identifiers: Gene Symbol (e.g., FOXP2), UCSC ID (e.g., uc003wys.3), RefSeq ID (e.g., NM_000669), NCBI Entrez ID (e.g., 7051), Ensembl Gene ID (e.g., ENSG00000117054) Ensembl Transcript ID (ENST00000008440) and UniGene ID (e.g., Hs.104894).

### Statistical analyses

Looking for enrichment or depletion within a gene list necessitates the use of a background list against which all comparisons are carried out. In principle, there are two possible ways to compile such default background lists. The first is taking all the known genes of the species. The second is taking only the subset of genes for which the DB contains annotations, namely, all genes that are linked to a phenotype. As described by Huang *et al*.^2^, the latter represents a more conservative approach that minimizes potential biases and thus it was the method of choice for Gene ORGANizer. This method assures that even if certain anatomical parts are over-represented in the ontology (as some phenotypes are easier to detect, or some diseases are more studied), using the entire list of annotated genes as background, instead of the whole genome, eliminates any bias towards them.

Users can choose whether to treat multiple transcripts of the same gene as one record (*gene-based* analysis) or as separate records (t*ranscript-based* analysis). Whereas gene-based analysis would fit most applications, transcript-based analysis can be useful when there is a biological meaning for two different transcripts appearing in the gene list.

Let *N_i_* be the number of genes associated with organ in the background list. Of them, suppose that *n_i_* ≤ *N_i_* genes are present in our input gene list. Let *N* be the total number of background genes, of which a total of *n* appear in our input list. The significance level of the enrichment or depletion is computed using the hypergeometric distribution *h*(*n_i_*; *N*, *N_i_*, *n*), where *P*-values are computed using the mid-range correction^54^. The user can choose to correct multiple comparisons through either the Bonferroni correction or FDR. Naturally, there is some degree of correlation between the different body parts. For example, genes that affect the small intestine, will often affect the large intestine as well. As described by Zhang et al.^55^, such correlations only make the computed FDR more conservative. Enrichment or depletion of organ is reported as

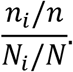

### Controls and validation

In order to test whether Gene ORGANizer identifies enrichment of specific body parts in lists where such enrichments are expected, or known to exist, we ran in Gene ORGANizer several gene lists, detailed in the main text. Unless otherwise stated, in all analyses we used *confident* and *typical+non-typical* associations, as they represent the default, most curated and most useful options in Gene ORGANizer.

Running the endocrine-related genes on KEGG in Gene ORGANizer revealed the ubiquitous effects of the endocrine system, with 67 organs significantly enriched, the top ones being lymphatic organs, endocrine glands and reproductive organs. Within the nervous system-related genes, we found enrichment in the brain and the cerebellum, although this trend is not significant (FDR = 0.689 for both). In fact, we detect no significantly enriched body parts, probably owing to the fact that most genes in the nervous and sensory categories in KEGG are involved in synapse biology– a basic function across all body parts. Within digestion-related genes, Gene ORGANizer detected enrichment of digestion-related organs (Supplementary Fig. S1).

---

**Supplementary Fig. 1.**
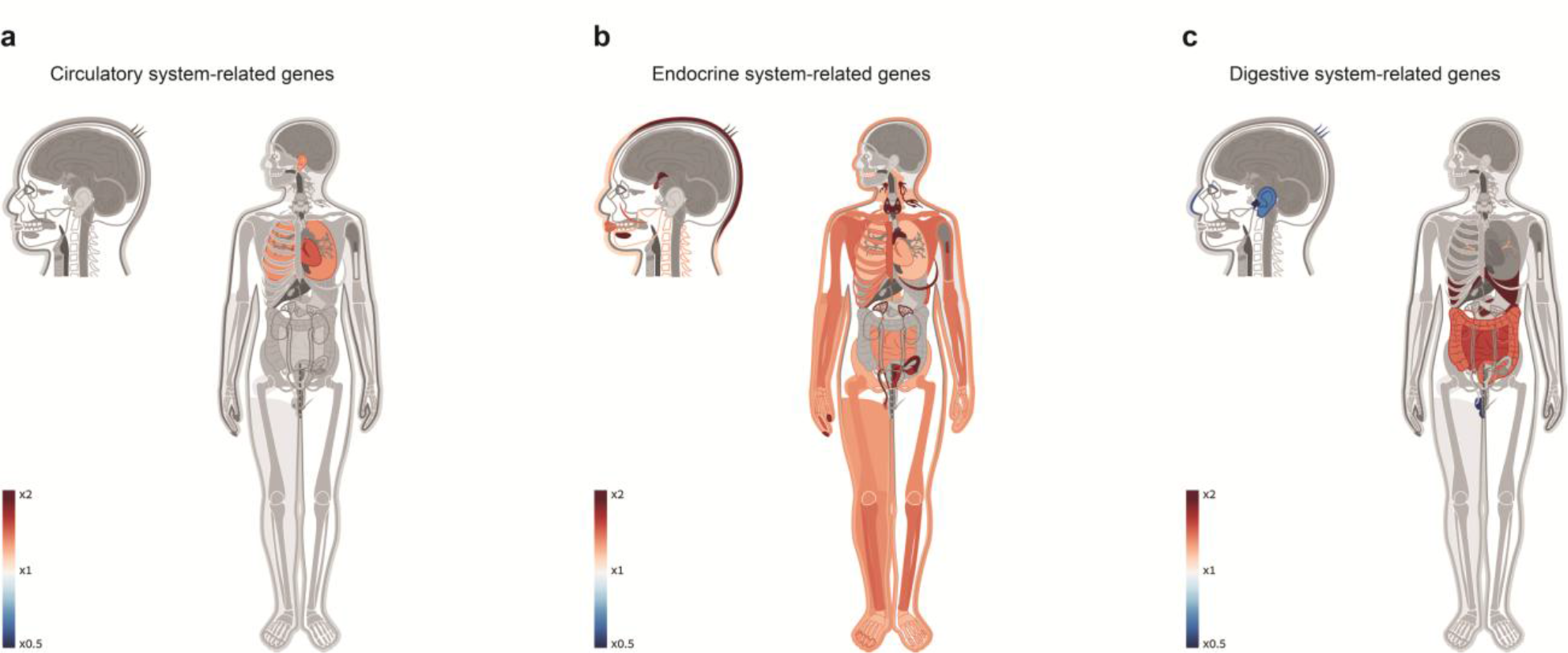
Gene ORGANizer correctly identifies the organs that are known to be regulated by a group of genes. **a.** Genes that are linked to the circulatory system in KEGG exhibit significant enrichment in circulation-related organs. **b.** Genes that are linked to the endocrine system exhibit significant enrichment in endocrine-related organs, as well as in many other organs, owing to their ubiquitous function. **c.** Genes that are linked to the digestive system exhibit significant enrichment in digestion related organs.

**Supplementary Fig. 2.**
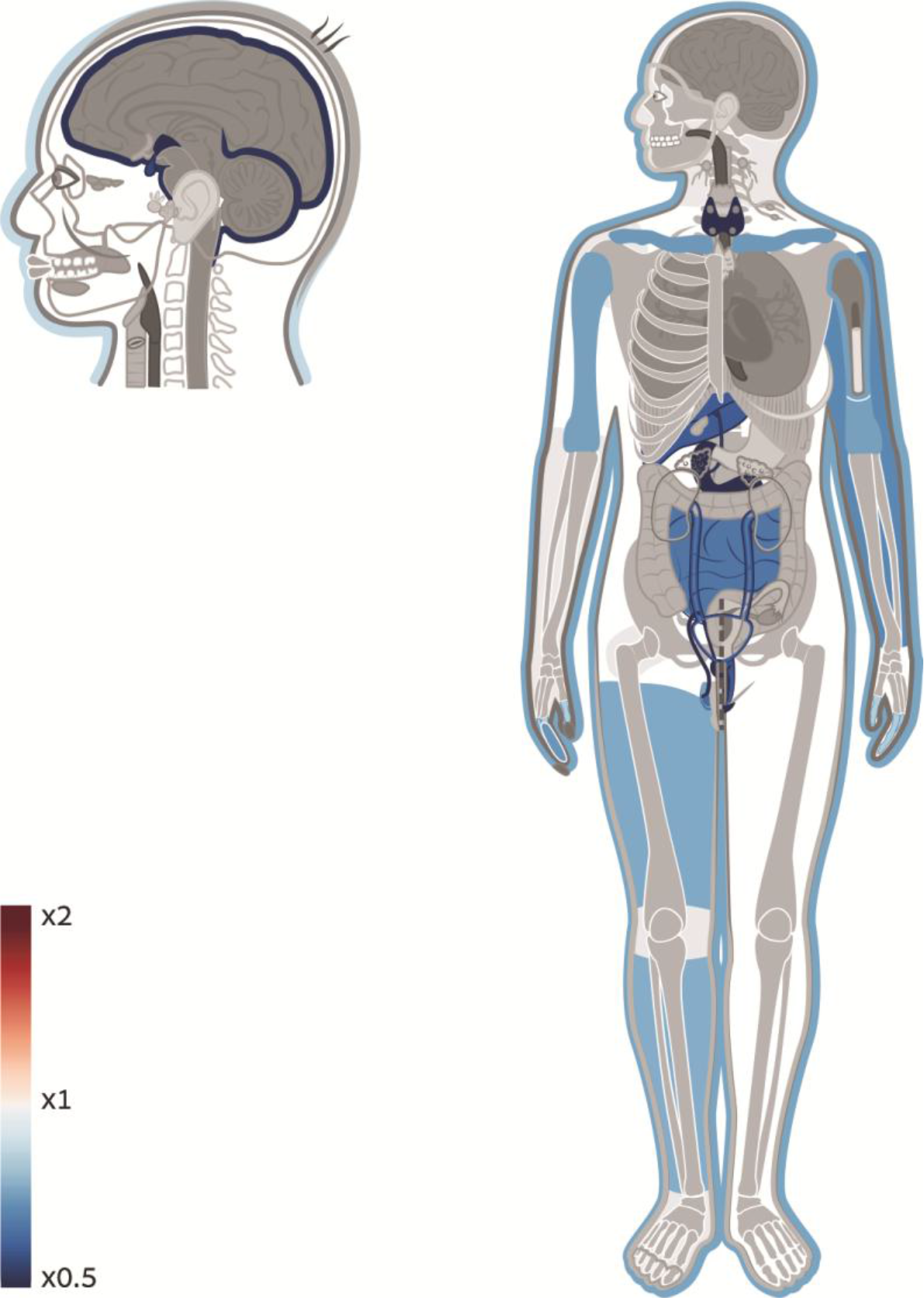
Candidate genes that went through positive selection in human and great ape lineages tend to affect less organs. A body and head map of enrichment and depletion of organs across genes where signatures of selective sweeps were detected along *Hominidae* lineages. 26 organs are significantly depleted, suggesting that positively selected genes have more constrained and organ-specific functions compared to the rest of the genome.

